# Effects of microplastics mixed with natural particles on *Daphnia magna* populations

**DOI:** 10.1101/2022.05.04.490562

**Authors:** Christoph Schür, Joana Beck, Scott Lambert, Christian Scherer, Jörg Oehlmann, Martin Wagner

## Abstract

The toxicity of microplastics on *Daphnia magna* as a key model for freshwater zooplankton is well described. While several studies predict population-level effects based on short-term, individual-level responses, only very few have validated these predictions experimentally. Thus, we exposed *D. magna* populations to irregular polystyrene microplastics and diatomite as natural particle (both ≤ 63 μm) over 50 days. We used mixtures of both particle types at fixed particle concentrations (50,000 particles mL^-1^) and recorded the effects on overall population size and structure, the size of the individual animals, and resting egg production. Particle exposure adversely affected the population size and structure and induced resting egg production. The terminal population size was 28–42% lower in exposed compared to control populations. Interestingly, mixtures containing diatomite induced stronger effects than microplastics alone, highlighting that natural particles are not *per se* less toxic than microplastics. Our results demonstrate that an exposure to synthetic and natural particles has negative population-level effects on zooplankton. Understanding the mixture toxicity of microplastics and natural particles is important given that aquatic organisms will experience exposure to both. Just as for chemical pollutants, better knowledge of such joint effects is essential to fully understand the environmental impacts of complex particle mixtures.

**Environmental Implications:** While microplastics are commonly considered hazardous based on individual-level effects, there is a dearth of information on how they affect populations. Since the latter is key for understanding the environmental impacts of microplastics, we investigated how particle exposures affect the population size and structure of *Daphnia magna*. In addition, we used mixtures of microplastics and natural particles because neither occurs alone in nature and joint effects can be expected in an environmentally realistic scenario. We show that such mixtures adversely affect daphnid populations and highlight that population-level and mixture-toxicity designs are one important step towards more environmental realism in microplastics research.

**Graphical Abstract:** 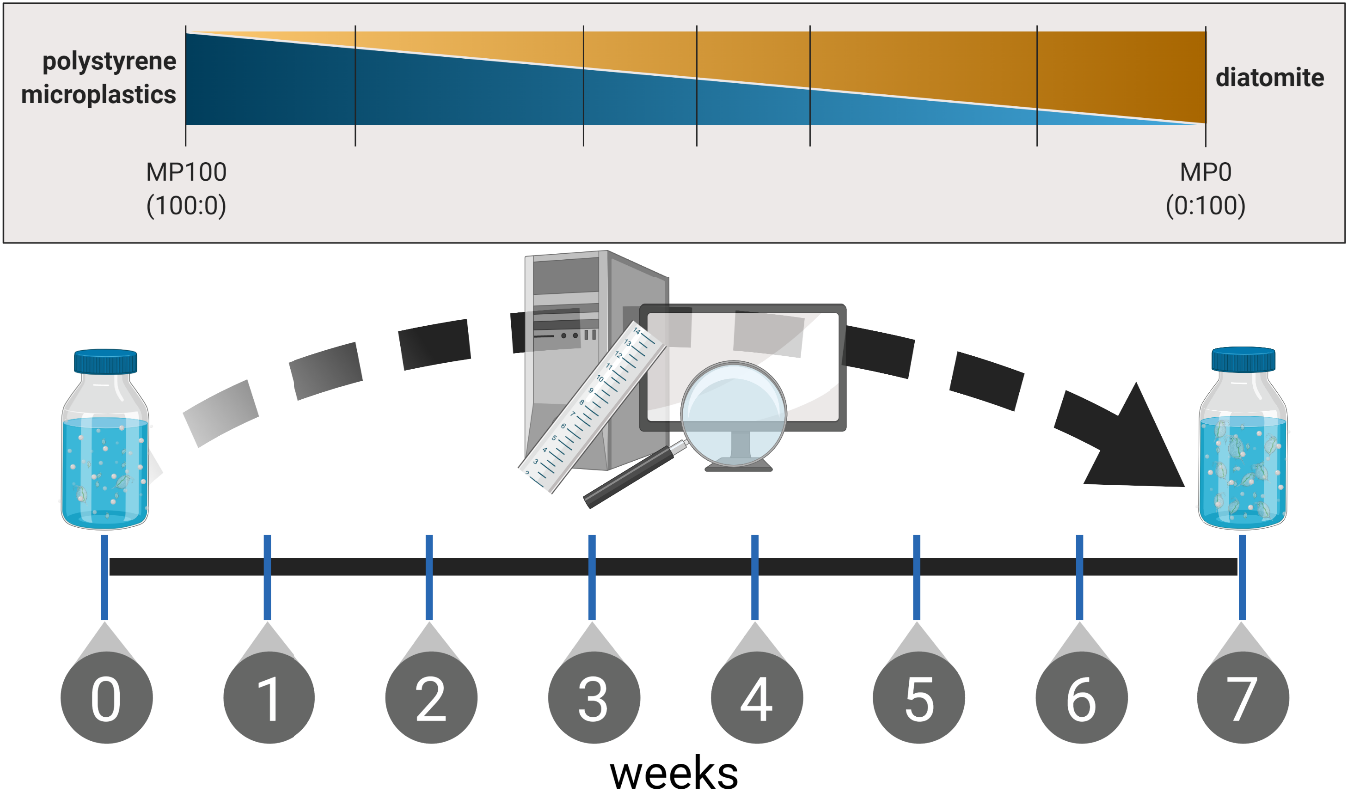

**Highlights:**

- *Daphnia* populations exposed to mixtures of microplastics and diatomite
- Effects on population size, structure, and resting egg production
- Diatomite as natural particle more toxic than microplastics
- Particle mixtures induce negative population-level effects
- Particle mixtures represent more realistic exposure scenario

## Introduction

Microplastics (MP) are ubiquitous pollutants in the aquatic environment (Čerkasova et al. 2023). They can interact with and affect a broad range of species across all levels of biological organization (Triebskorn et al. 2018). Indeed, meta-analyses using species-sensitivity distributions indicate that adverse effects in aquatic species occur at relatively low concentrations typically found in polluted habitats. For instance, Mehinto et al. determined that ambient concentrations of 0.3–5 MP/L (50–900 μm/L) would put 5% of the most sensitive species at risk (Mehinto et al. 2022). This is consistent with other studies (Adam, Von Wyl, and Nowack 2021; Yang and Nowack 2020; Adam, Yang, and Nowack 2018; Takeshita et al. 2022). MPs affect aquatic organisms at all levels of biological organization, ranging from molecular, cellular, and tissue-level effects to impacts on the life history of individuals. Here, numerous reviews are available on aquatic species in general (Rezania et al. 2018; Du et al. 2021; Wang et al. 2019; Elizalde-Velázquez and Gómez-Oliván 2021; Huang et al. 2021; Triebskorn et al. 2018) and zooplankton in particular (Foley et al. 2018; Botterell et al. 2019; Yin et al. 2023). Regarding the latter, the Cladoceran *Daphnia magna* is one of the most studied species due to it ecological role as keystone species and widespread establishment as ecotoxicological model (Castro-Castellon et al. 2022).

Importantly, the current literature on the toxicity of MP in Daphnia species is strongly biased towards acute exposure scenarios, that is, animals being exposed to MP for very short periods, only. This is despite the fact that, due to their short generation time and the environmental persistence of MP, daphnids are exposed continuously over generations and not just intermittently (Rozman and Kalčikova 2021; Yin et al. 2023). Daphnids as r-strategists form large, often short-lived, populations. Population growth rates are high, but quickly reach a carrying capacity limited by space and/or food. Such stressors are then often met with the formation of resting eggs that can resurrect the population once conditions have returned to a more favorable state (Smirnov 2017). Contrary to that, only little information is available on the long-term, multi-generational, or population-level effects of MP on Daphnia species (Yin et al. 2023; Junaid et al. 2023). Another bias in experimental studies is the use of spherical MP that are not representative of particles that are most abundant in the environment, especially plastic fibers and irregular plastic fragments (E. E. Burns and Boxall 2018; Rozman and Kalčikova 2021). Thus, a long-term, continuous exposure to irregularly-shaped MP throughout an individual’s lifetime, as well as following generations, is a more realistic scenario (Schür et al. 2020; 2021).

In the environment, organisms interact with all kinds of natural, non-food particles in addition to MP, that is, suspended solids consisting of fine organic and inorganic matter typically < 62 μm (Bilotta and Brazier 2008). These naturally occurring particles negatively affect daphnids, sometimes across generations (Kirk 1991b; 1991a; Robinson, Capper, and Klaine 2010; Ogonowski et al. 2016; Schür et al. 2020). Nonetheless, as non-selectively filter-feeding organisms, daphnids are well-adapted to non-food particles. This is achieved through a number of behavioral and physiological mechanisms, including a reduction in feeding rate, regurgitation of boluses, and the ability to remove adhering particles from the filtering setae via the post-abdominal claw (C. W. Burns 1968a; 1968b; Kirk 1991a; Ogonowski et al. 2016). Since exposure in the environment is never to a single type of particle (synthetic or natural) and the effects of suspended solids in comparison to MP are often overlooked but important to benchmark the toxicity of MP (Scherer et al. 2018). Indeed, two recent meta-analyses show that MP are more toxic than suspended solids but, at the same time, highlight the dearth of comparative studies and inconsistencies in methodology (M. Ogonowski et al. 2022; Doyle, Sundh, and Almroth 2022). Thus, exposing animals to both MP and natural particles, alone and in mixture, is important to increase our understanding of particle toxicity in a more environmentally relevant setting (Gerdes et al. 2018; 2019).

Since there is little knowledge on the population-level effects of MP in comparison to other suspended solids, the aim of this study is to quantify the effects of irregular polystyrene MP and of diatomite as a natural particle on *D. magna* populations. We designed an experiment in which daphnid populations with a defined age structure and size were continuously exposed over 50 d to MP and diatomite, either alone or in mixtures, using a constant particle number of 50,000 particles mL^-1^ and constant food levels. These concentrations were not intended to reflect the environmental levels of plastics or natural particles. They rather serve as a proof of concept for benchmarking the effects of MP to natural particles as well as for investigating the impacts of particle mixtures on populations in a more realistic scenario.

## Materials and methods

### Daphnia culture

Ten *D. magna* individuals were cultured in 1 L of Elendt M4 medium (Elendt and Bias 1990; OECD 2012) at 20 °C with a 16:8 h light:dark cycle. The daphnids were fed with the green algae *Desmodesmus subspicatus*. Algae suspension is was added to the culture vessels thrice a week to achieve a mean food level of 0.2 mg carbon per individual per day (mgC daphnid^− 1^ d^− 1^). The medium was fully renewed once a week.

### Particle preparation

The irregularly shaped MP were produced from polystyrene coffee-to-go-cup lids obtained from a local bakery. They were rinsed, cut into pieces using scissors, frozen in liquid nitrogen and ground up in a ballmill (Retsch MM400, Retsch GmbH, Germany) at 30 Hz for 30 s. Diatomite (SiO_2_) was purchased from Sigma Aldrich (CAS: 91053-39-3). Both particle types were sieved to ≤ 63 μm using a sediment shaker (Retsch AS 200 basic, Retsch GmbH, Germany) to achieve particles in a size range that daphnids can ingest (0.2–75 μm; Scherer et al. (2018)). Particle size distributions within the measuring margins of the Coulter counter (Multisizer 3, Beckman Coulter, Germany; orifice tube with 100 mm aperture diameter for a particle size range of 2.0–60 mm; measurements in filtrated (< 0.2 μm) 0.98% NaCl solution) are given in the supplementary material (Figure S1). Size distributions for the diatomite size fraction < 2 μm for a suspension prepared with the same method are given in the supplementary materials of Scherer et al. (2019)) but were not measured for this study. Furthermore, Scanning electron microscope images of both particle types were taken using a Hitachi S-4500 scanning electron microscope (supplementary material, Figure S2). For that, 20 μL of each suspension was transferred to the sample holder, dried under a heat lamp, and sputtered with gold before imaging. Additional characterization of similar materials and particle types can be found in Schür et al. (2021) and Scherer et al. (2019). In terms of chemicals composition, the MP contained the chemicals present in the product (i.e., intentionally and non-intentionally added substances) and the diatomite was purified and contained ≤ 1% of HCl-soluble matter according to the manufacturer.

Exposure suspensions were prepared by dilution in Elendt M4 medium based on measured particle concentrations of the stock solutions (Multisizer 3, Beckman Coulter) and used throughout the experiment. Previous experiments, described in Schür et al. (2020), showed a good correlation between nominal and measured particle concentrations. A new MP stock suspension was prepared after day 37. Fourier transform infrared spectroscopy (ATR-FTIR) spectra (FTIR Spectrum Two, PerkinElmer; LiTa03 detector, range: 4000−450 cm^-1^) of the raw plastic material before and after grinding and sieving are given in Figure S3 of the supplementary material.

### Experimental design

The initial daphnid populations consisted of 3 adults (2 weeks old), 5 juveniles (1 week old), and 8 neonates (< 72 h old) held in 1 L glass vessels containing 900 mL Elendt M4 medium (OECD 2012). This composition was chosen to start with a population structure that represented all age groups and to ensure an early onset of reproduction. Each population was kept for 50 d and fed a constant ration of 0.5 mgC d^-1^ of *D. subspicatus*. All treatment groups were exposed to a total of 50,000 particles mL^-1^ of varying ratios of MP and diatomite (n = 3, Table 1).

**Table 1:** Ratios and absolute nominal particle concentrations of microplastics and diatomite in the treatment groups of the population experiment.

| Treatment group | Microplastics |  | Diatomite |  |
| --- | --- | --- | --- | --- |
|  | % | Particles mL<sup>-1</sup> | % | Particles mL<sup>-1</sup> |
| Control | 0 | 0 | 0 | 0 |
| MP100 | 100 | 50,000 | 0 | 0 |
| MP80 | 80 | 40,000 | 20 | 10,000 |
| MP60 | 60 | 30,000 | 40 | 20,000 |
| **MP50** | 50 | 25,000 | 50 | 25,000 |
| **MP40** | 40 | 20,000 | 60 | 30,000 |
| **MP20** | 20 | 10,000 | 80 | 40,000 |
| **MP0** | 0 | 0 | 100 | 50,000 |

Populations were fed thrice per week, and the medium was fully exchanged with new medium or particle suspensions on days 7, 14, 21, 28, 37, 42, and 50. During each feeding, vessels were covered with a lid and gently inverted to re-suspend the particles. With each medium exchange, populations were sieved, transferred to an hourglass, and photographed. ImageJ (Schindelin et al. 2012) was used to then quantify living animals (total population size, Figure 1, Figure S4), the number of resting eggs (Figure 2, Figure S5), body lengths (Figure 3, Figure S6). Resting eggs are seen as indicators of population stress like insufficient food or high population density (Smirnov 2017). Individual body lengths were measured from the center of the eye to the base of the apical spinus (Ogonowski et al. 2016). Body lengths were categorized into three size/age classes in accordance with Agatz et al. (2015), including neonates (≤ 1400 μm), juveniles (1400–2600 μm), and adults (> 2600 μm).

**Figure 1:**
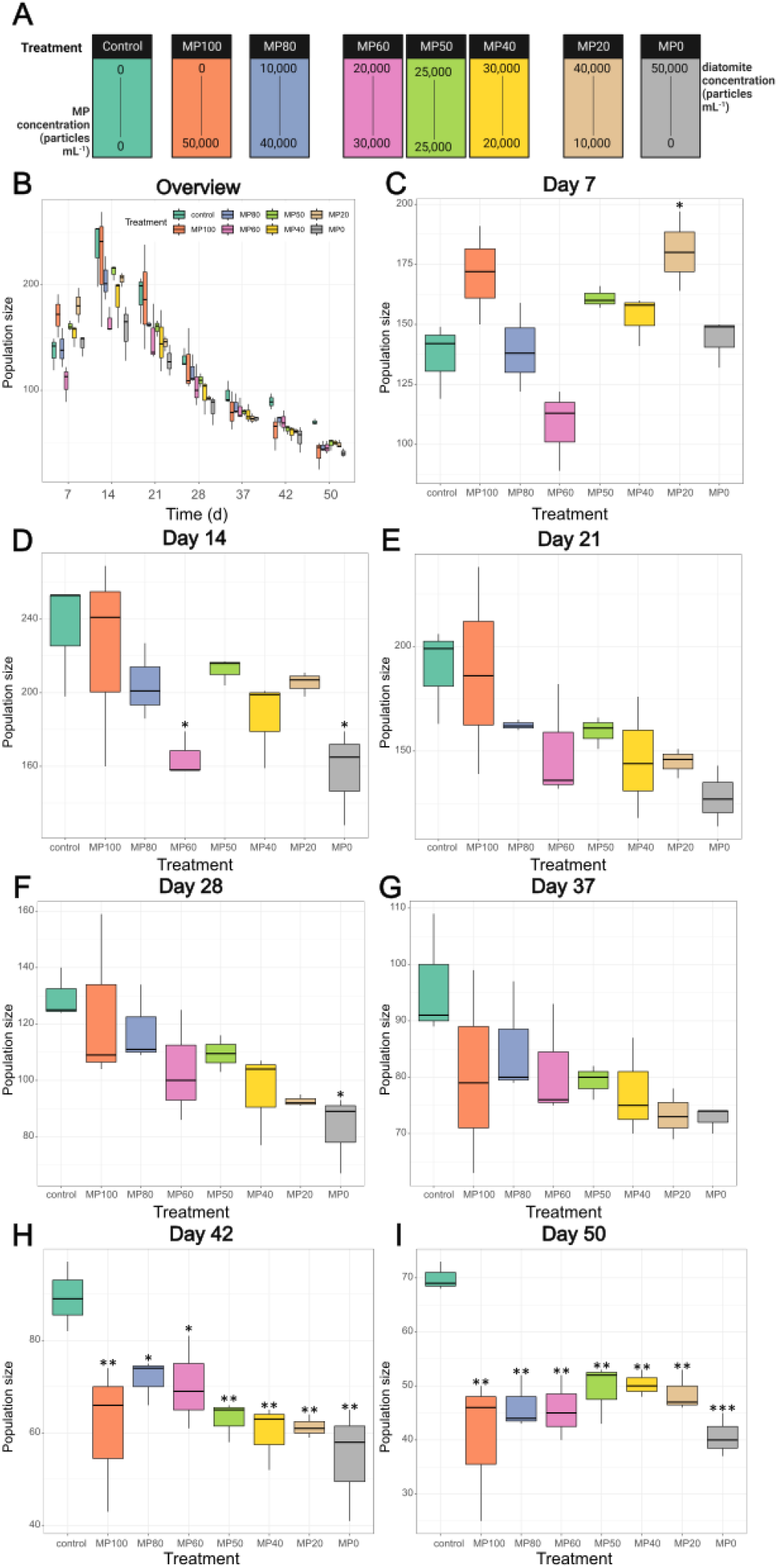
Boxplots of the population size of *Daphnia magna* exposed to polystyrene microplastics (MP100), diatomite (MP0), or their mixtures over 50 d (B). C–I represent the population sizes after 7, 14, 21, 28, 37, 42 and 50 d, respectively. Significant differences compared to control populations are indicated by asterisks: * p < 0.05, ** p < 0.01, *** p < 0.001. Subfigure A indicates the concentrations of the two particle types in the treatments.

**Figure 2:**
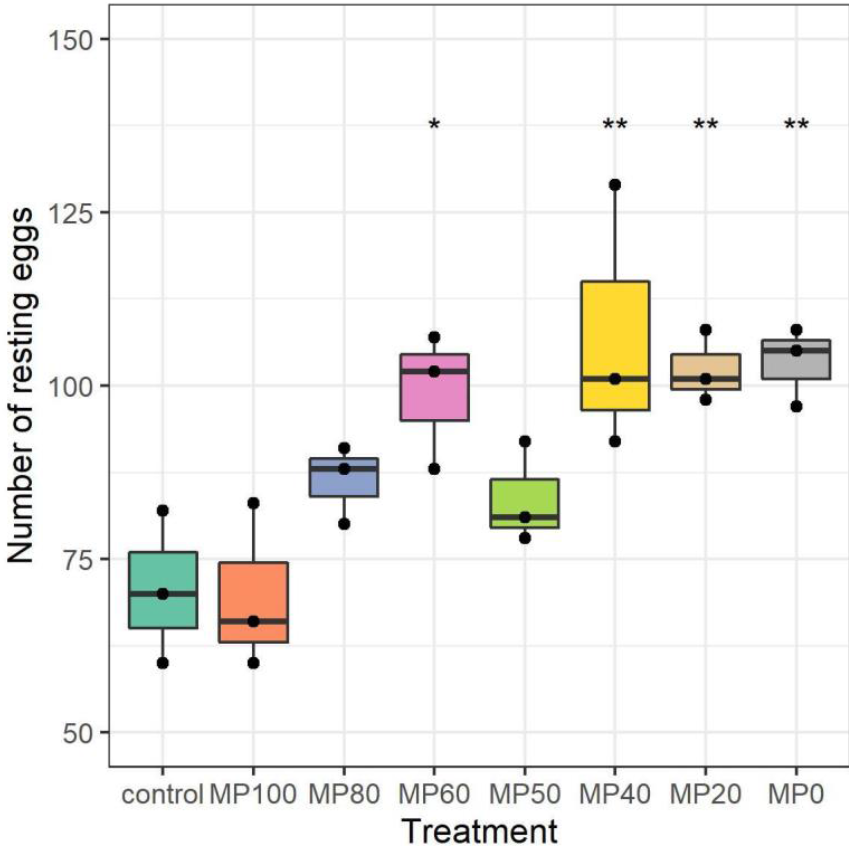
Total number of resting eggs produced by *Daphnia magna* populations exposed to polystyrene microplastics (MP100), diatomite (MP0), or their mixtures over 50 d (n = 3). Significant differences compared to control populations are indicated by asterisks: * p < 0.05, ** p < 0.01.

**Figure 3:**
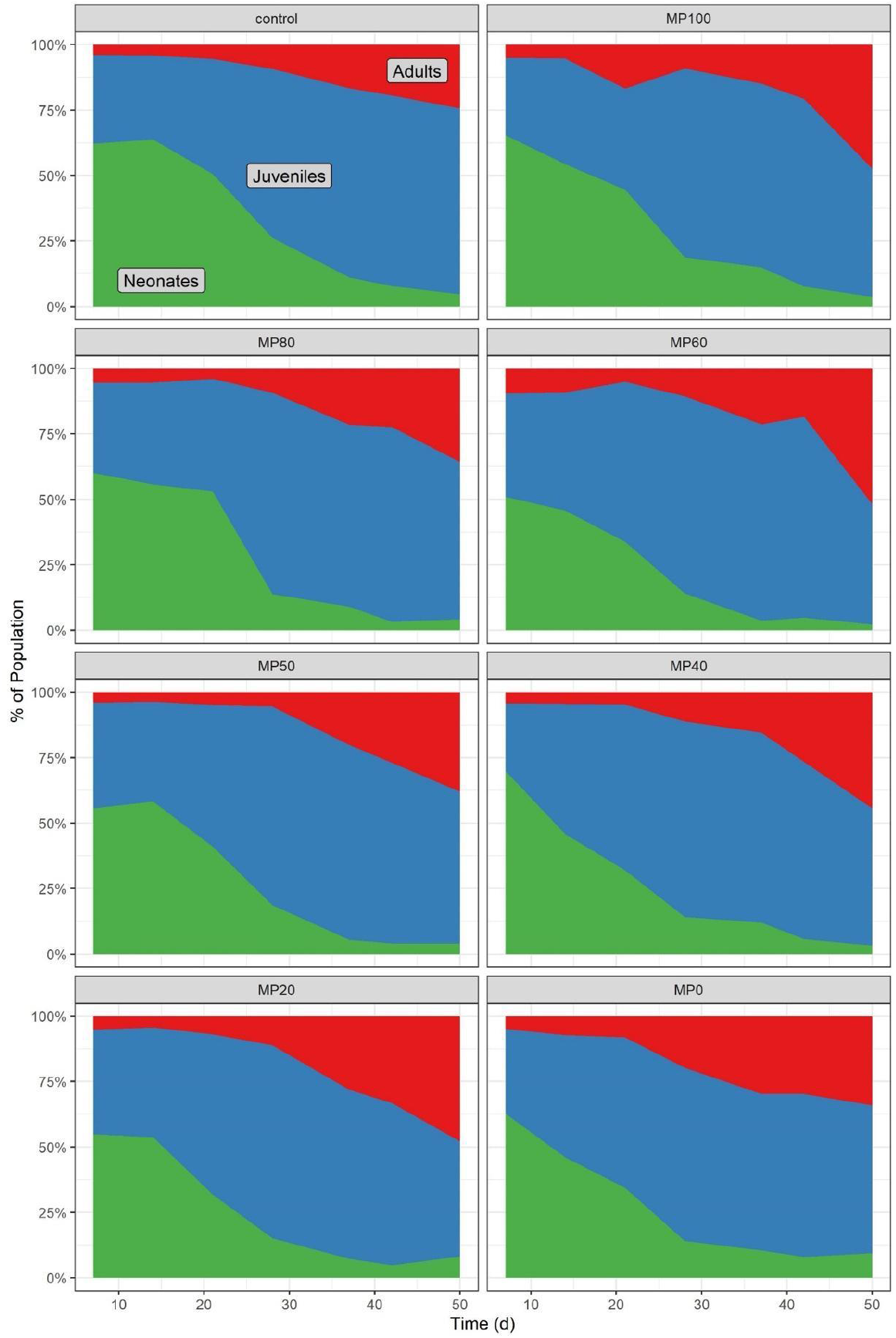
Population structure of *Daphnia magna* exposed to polystyrene microplastics (MP100), diatomite (MP0), or their mixtures over 50 d. Data presented as mean relative ratios of neonates (green), juveniles (blue), and adults (red) compared to the overall population size (n = 3).

### Statistical analysis

The data was visualized using R (R Core Team 2021) with RStudio 2021.09.2+382 and the tidyverse package (Wickham et al. 2019). The impact of exposure time and treatment on population sizes (i.e., total number of animals at a given time) and structure was analyzed using a one-way ANOVA with Holm-Šídák’s multiple comparison test against the corresponding control group for each time point in GraphPad Prism for Mac 9.3.1. The number of resting eggs on day 50 of the experiment was compared against the control group using a one-way ANOVA with Holm-Šídák’s multiple comparisons test in GraphPad Prism for Mac 9.3.1. The body length of individuals in each population was compared using Kruskal-Wallis tests followed by Dunn’s multiple comparison tests. Boxplots are created with the geom_boxplot() function of the ggplot2 package (Wickham 2016) in accordance with Mcgill et al. (1978). Significance levels are indicated by asterisks as follows: * p < 0.05, ** p < 0.01, *** p < 0.001.

## Results

Following the goal to investigate the population-level effects of mixtures of MP and natural particles, the experiment included three main endpoints: population size (i.e., total number of individuals per population at each time point), population structure based on the body lengths of the individuals comprising each population, and the number of resting eggs (*ephippiae*) per population.

### Population size

All populations, both exposed to particles and of the control group, grew rapidly with regards to the number of individuals during the first two weeks, with little variability between the three replicates per treatment group (Figure 1). We hypothesize that this is because the available food was sufficient for such small populations coupled with low initial population densities acting as triggers for rapid population growth. All population sizes peaked at day 14, declined from day 21 onwards, and reached their lowest size on day 50.

We observed a concentration-dependent effect in the populations exposed to particles in such that in the phase of rapid decline (days 21–37), daphnid populations exposed to particle mixtures that contained more diatomite had a lower population size (Figure 1, Figure S4). For instance, populations exposed to 100% diatomite (MP0) consisted of significantly fewer animals than the control populations on day 14 and 28 (p < 0.05, one-way ANOVA). At the end of the experiment, on day 42 and 50, the size of all populations exposed to MP, diatomite or mixtures thereof was significantly smaller than the control populations (p < 0.05). Notably, the terminal population size in all treatments was 28–42% lower compared to control.

### Resting egg formation

Resting egg formation occurred in all populations, including controls, after day 14 but to varying degrees (Figure S5). Since the production of resting eggs is a stress response (Smirnov 2017), this indicates a rapid onset of stress caused by increasing population densities and/or decreasing food levels. The particle exposure had a significant effect on the total number of resting eggs produced, with the populations in the MP60 (p = 0.023), MP40 (p = 0.003), MP20 (p = 0.011), and MP0 (p = 0.008) groups producing circa 100 ephippiae compared to 70 in the control populations (Figure 2). Similar to the effect on intermediate population sizes, this points towards a stronger effect of diatomite compared to MP.

### Individual body length

We measured the body length of each individual in a population weekly and used this to describe the population structure by categorizing the daphnids into neonates, juveniles, and adults. The initial population growth is largely driven by the production of neonates (Figure 3). As a result of the lower reproduction from day 14 onwards, the population structure shifts towards juveniles and adults. Particle exposure affected the number of neonates and juveniles in the populations (Tables S2 and S4): At day 28, populations exposed to diatomite or diatomite mixed with 20, 40, 60 and 80% MP consisted of significantly fewer neonates. This effect translated into significantly fewer juveniles in all particle-exposed populations on day 50. The number of adults was low compared to the other size classes and increased over the course of the experiment without significant impact of the particle exposure. In accordance with the loss of neonates and juveniles, individuals in particle-exposed populations were in many cases significantly larger than in control populations (Kruskal-Wallis tests, Table S6).

## Discussion

We exposed *D. magna* populations to 50,000 particles mL^-1^ of either polystyrene MP, diatomite, or mixtures of both over the course of 50 d. Notwithstanding the ratio of MP to diatomite, particle exposure significantly reduced the population size from day 43 onwards and resulted in populations consisting of 28–42% less individuals than control populations at the end of the experiment. This effect on population size is most likely due to particle exposures having a negative impact on reproduction as previously shown by Ogonowski et al. (2016) and Schür et al. (2020), especially after the population size has peaked. The reproductive toxicity of particles is also reflected in the population structure with particle-exposed populations consisting of larger and, thus, older individuals than control populations. In addition, a reduced availability of food contributes to limiting the population growth over the course of the experiment, in such that population growth results in less food being available per individual (Figure 1B). While this affects all treatment groups, less food is available to control individuals (e.g., 5.6 μgC d^-1^ invividual^-1^ at 42 d) than to particle exposed daphnids (7.9 μgC d^-1^ invividual^-1^ at 42 d). Thus, the food limitation did not mask the effect of the particle exposure. Taken together, this demonstrates that mixtures of synthetic and natural particles have negative effects at the population level in *D. magna*.

The fact that MP as well as their mixtures with natural particles affected the population size and structure highlights that the well-documented individual-level toxicity of MP and other particles in daphnids translates into impacts at the population level. While multiple studies predict effects of MP exposures on population growth rates based on individual level responses (e.g., Martins and Guilhermino (2018); Guilhermino et al. (2021)), population and multigeneration effects were recently identified as severe data gaps in a review on the ecotoxicology of MP in Daphnia (Yin et al. 2023). Bosker et al. (2019) reported that exposure to polystyrene beads caused a significant decline in population size and biomass but did not affect the size of individuals or *ephippiae* production. Besides using another type of MP, their general approach was different from ours as they grew populations to holding capacity before starting particle exposure at day 30. This probably reduced the overall stress level induced by continuous particle exposures.

Zebrowski et al. (2022) investigated how the exposure to polystyrene, high-density polyethylene (PE), and the assumed biodegradable polyhydroxybutyrate (PHB) affected the growth (measured by population density, i.e., individuals per L) of competing populations of *D. magna, Daphnia pulex*, and *Daphnia galeata* under constant food levels (two species were paired in competition). While the outcome of the competition experiments was not affected (the same species outperformed their competitors as in the particle-free control treatments), two main findings are worth mentioning: (I) The larger *D. magna* did not always outcompete the smaller cladoceran species, but only the smallest *D. galeata*, while *D. pulex* consistently persisted against both other species, and (II) *D. magna* and *D. pulex* populations were affected by polystyrene particle exposure, compared to the respective control groups, while *D. galeata* populations grew very similarly to their control populations. Some of their results hint towards a positive effect of the PHB exposure, about which the authors hypothesize that the material is biodegraded and serves as nutrients for bacteria in the exposure vessel, which subsequently improve the nutritional status of the cladcoerans.

Al-Jaibachi et al. (2019) observed the initial decline but subsequent recovery of daphnid populations in MP-exposed mesocosms, while no effect on other species was observed. Here, high variability and unknown influencing factors from the mesocosm setup impede the comparison between the two studies. Nonetheless, all three studies demonstrate that MP effects also manifest on the population level, which is considered highly relevant for assessing the environmental risks of these particles.

We used multiple mixtures of MP and diatomite at a fixed numerical concentration to explore a more realistic exposure scenario (i.e., MP as part of a more diverse set of suspended solids) and to investigate whether the mixtures’ toxicity is driven by plastic or natural particles. Indeed, our results show that diatomite is more toxic to daphnid populations than MP. With regards to the intermediate population size, resting egg production, and population structure, exposure to pure diatomite induced stronger effects than to pure MP (Figures 1–3). In the treatments with particle mixtures, we often observed a concentration-dependent response with mixtures containing more diatomite being more toxic. This is particularly obvious for the population sizes at days 14–28 and the resting egg production. Accordingly, particle mixtures consisting of more diatomite are more toxic.

The reason for the higher toxicity of diatomite compared to MP may be its porous and spiky structure (Figure S2). Diatomite has biocidal properties (European Food Safety Authority (EFSA) et al. 2020) and its absorptive and abrasive capacities will damage insect cuticles (Korunic 1998) and may injure the digestive system (Scherer et al. 2019). Diatomite is more toxic than MP despite the fact that it has a higher density and is larger than polystyrene particles (2.36 g cm^-3^ according to the manufacturer) and, thus, sediments faster. While we did not investigate the fate or uptake of the particles in this study, a faster settlement probably results in a lower exposure of daphnids to diatomite compared to polystyrene particles. This highlights= the challenges of comparing different particle types in terms of aligning their properties (Scherer et al. 2019) and keeping their bioavailable concentrations constant.

Diatomite has been used as a natural reference material in previous MP studies. In the freshwater mollusks *Dreissena polymorpha* and *Lymnea stagnalis*, diatomite was in general not more toxic than polystyrene MP (Weber, Jeckel, et al. 2021; Weber, von Randow, et al. 2021) but induced a stronger effect on the antioxidant capacity in the former species (Weber, Jeckel, and Wagner 2020) at identical numerical concentrations. In *Chironomus riparius* larvae, diatomite was toxic but less so than polyvinyl chloride MP at identical mass-based concentrations (Scherer et al. 2019). Since one of the main mechanisms of its toxicity appears to be the desorption of waxes from the cuticle, arthropods, such as chironomids and daphnids, may be particularly sensitive to diatomite exposures.

Our study shows that some natural particles can be more toxic than a mixture of natural particles and MP or MP by themselves. Earlier work compared the effects of the natural particle kaolin with polystyrene MP similar to those used in this study in a multigenerational study with daphnids (Schür et al. 2020). There, we found that kaolin had no effect, while MP affected all recorded endpoints in a concentration-dependent manner with effects increasing over generations. This shows that transferring findings on one particle type to another is not straightforward and MP may be more toxic than some but not all natural particles. Particle shape may play an important role as diatomite is spiky and sharp compared to kaolin which is rather round. Thus, the toxicity of natural particles will depend on their individual set of physicochemical properties and cannot be easily generalized without a better mechanistic understanding (see Scherer et al. (2019) for an in-depth discussion).

Finally, our study was not designed to mimic environmental concentrations of MP or natural particles. Instead, our aim was to investigate the toxicity of mixtures of both, because this exposure scenario is more realistic compared to the use of only MP in toxicity studies. Given that in nature, aquatic organisms will most likely be exposed to natural and synthetic particulate matter concurrently, a better understanding of the joint toxicity is needed to develop realistic predictions of environmental impacts.

## Conclusions

Our study demonstrates that an exposure to MP and diatomite alone as well as in mixture has negative population-level effects in *D. magna*. This corroborates previous predictions based on individual-level responses. Our findings are relevant because adverse impacts on populations of a keystone zooplankton species will have ecological consequences. However, the fact that we used one very high particle concentration only calls for follow-up studies to generate concentration-response relationships. We used mixtures of plastic and the natural particle diatomite because we deem this exposure scenario more realistic and found that diatomite is more toxic than MP. This contradicts the common assumption that natural particles are benign and highlights that – just as with MP – the toxicity of a particle type depends on its individual set of physicochemical properties. This calls into question whether general comparisons, such as MP are more or less toxic than something else, are meaningful. It also highlights the challenge of finding an adequate reference particle when attempting to perform such comparisons. Finally, we believe that investigating the mixture toxicity of synthetic and natural particles is valuable given that aquatic organisms will experience exposure to both. Similar to chemical pollutants, better knowledge of such joint effects is essential to fully understand the environmental risks complex particle mixtures pose to aquatic species.

## Author contributions

Christoph Schür: Conceptualization, Data curation, Formal analysis, Investigation, Methodology, Validation, Visualization, Project administration, Writing -original draft, Writing -review & editing Joana Beck: Data curation, Investigation, Writing -review & editing

Scott Lambert: Conceptualization, Methodology, Writing -review & editing

Christian Scherer: Conceptualization, Methodology, Investigation, Writing -review & editing

Jörg Oehlmann: Funding acquisition, Project administration, Resources, Writing -review & editing

Martin Wagner: Conceptualization, Formal analysis, Funding acquisition, Resources, Project administration, Visualization, Resources, Writing -review & editing

## Acknowledgments

This study was supported by the German Federal Ministry for Education and Research to CS, JO, and MW (02WRS1378I, 03F0789D). The graphical abstract was created with BioRender.

## Supplementary Material

The supplemental data are available ###.

