## Supplementary Material for "Effects of microplastics mixed with natural particles on *Daphnia magna* populations"

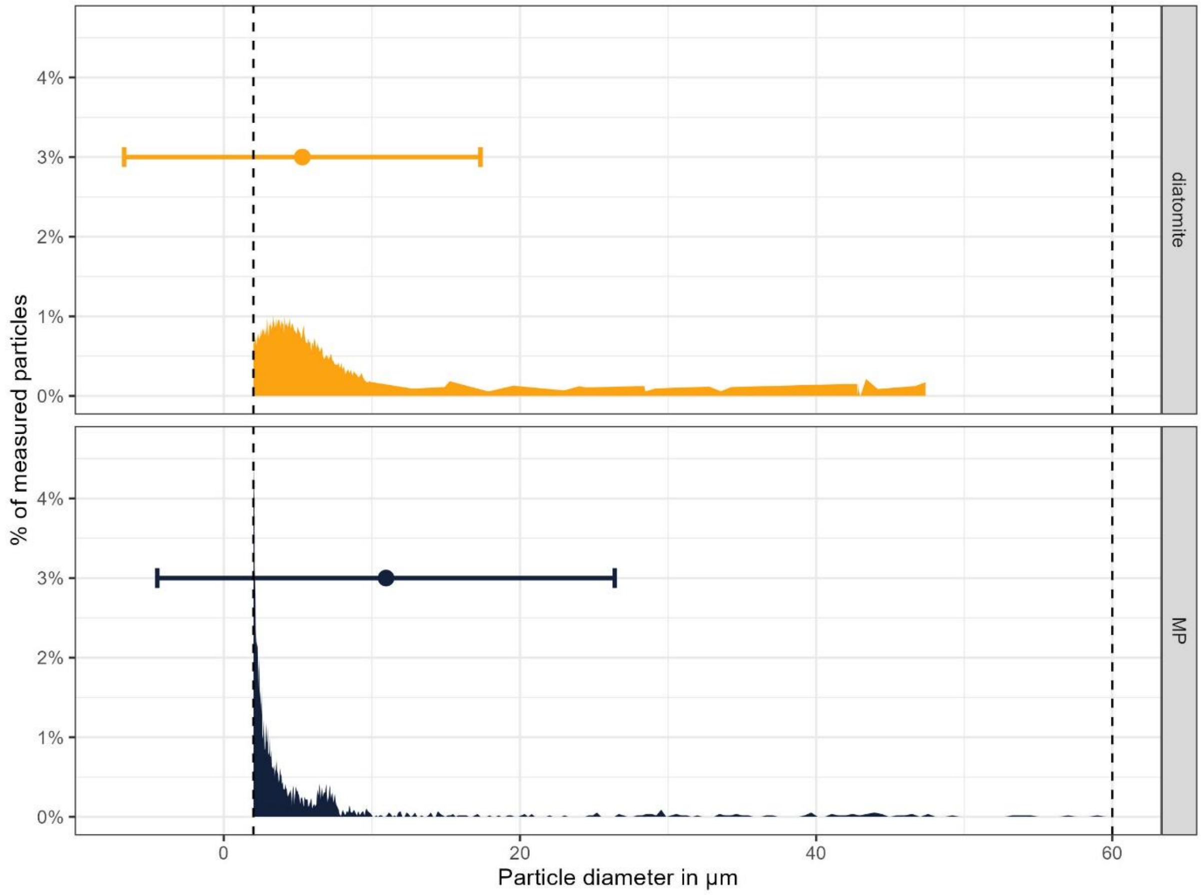

**Figure S1: Particle size distributions for diatomite (top, yellow) and MP (bottom, blue), measured in a Multisizer 3 for the size range ≥ 2 µm and ≤ 60 µm (dotted vertical lines).** The points and horizontal error bars at 3% indicate the median ± standard deviation (SD) of the particle sizes, respectively.

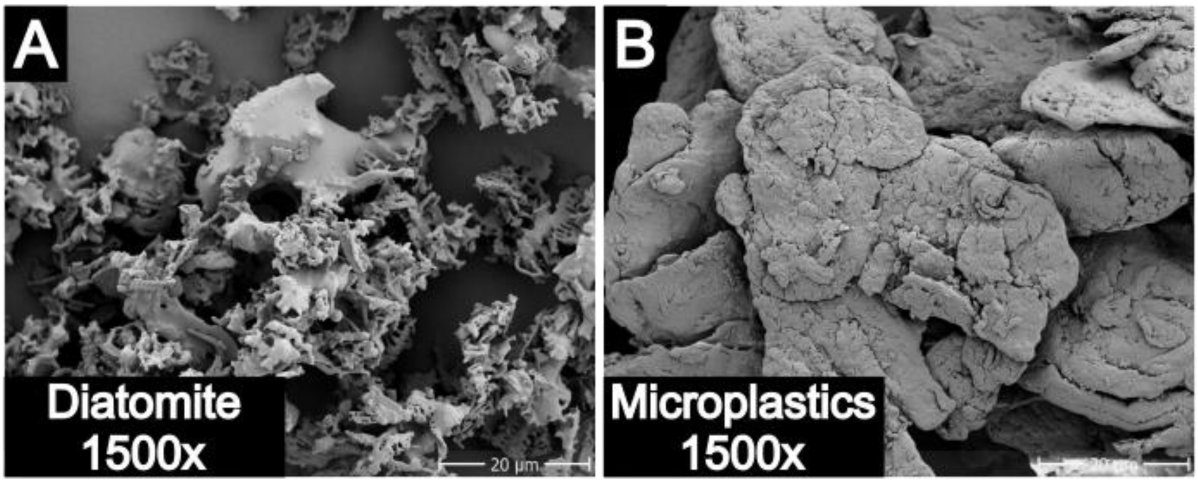

**Figure S2: Scanning electron microscopy micrographs of the two particle types (A: diatomite, B: polystyrene microplastics) used throughout this study (1500× magnification).** Image B was previously published as part of Figure 1 in Schür et al. (2021).

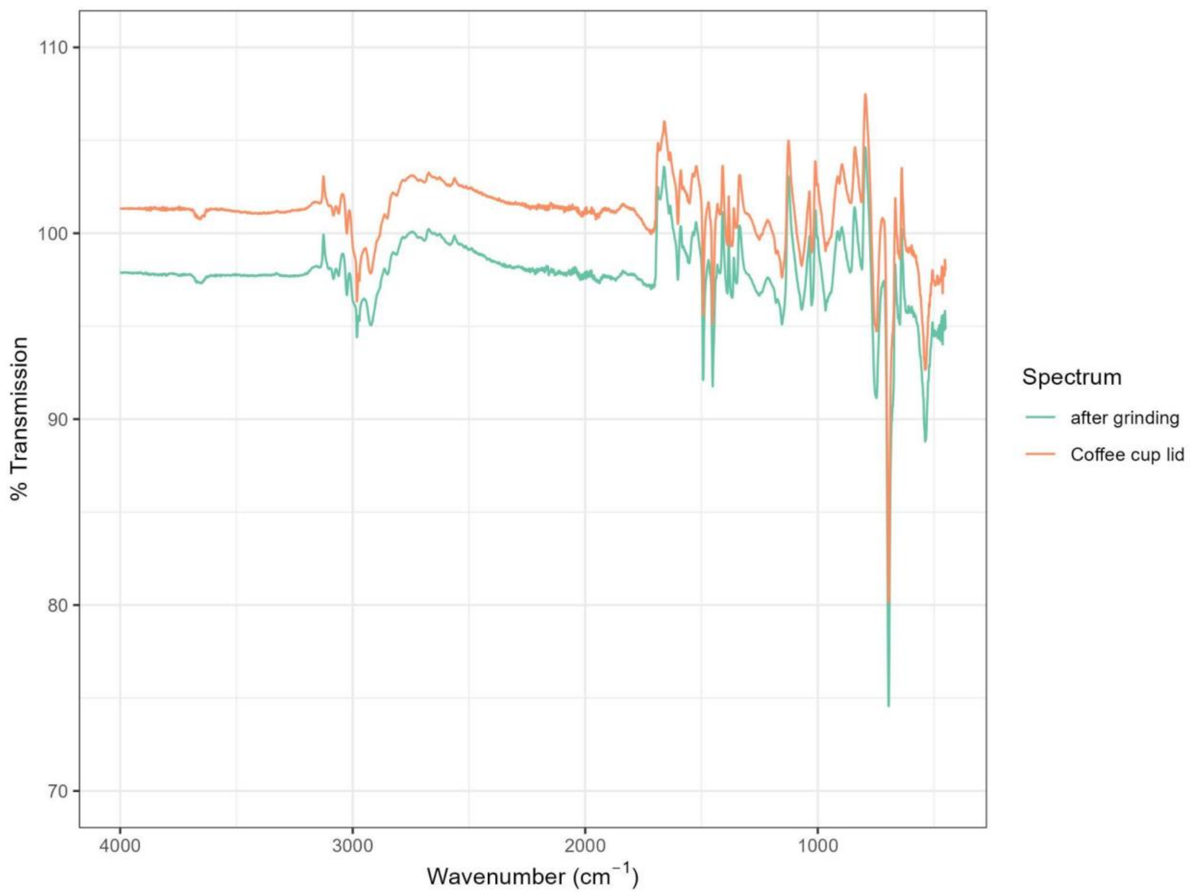

**Figure S3: Fourier-transform infrared spectroscopy (FTIR) spectra of the coffee-to-go cup lids that were the raw material for the microplastics in this study and the microplastics after grinding in the ballmill (see Materials and methods in the main manuscript).**

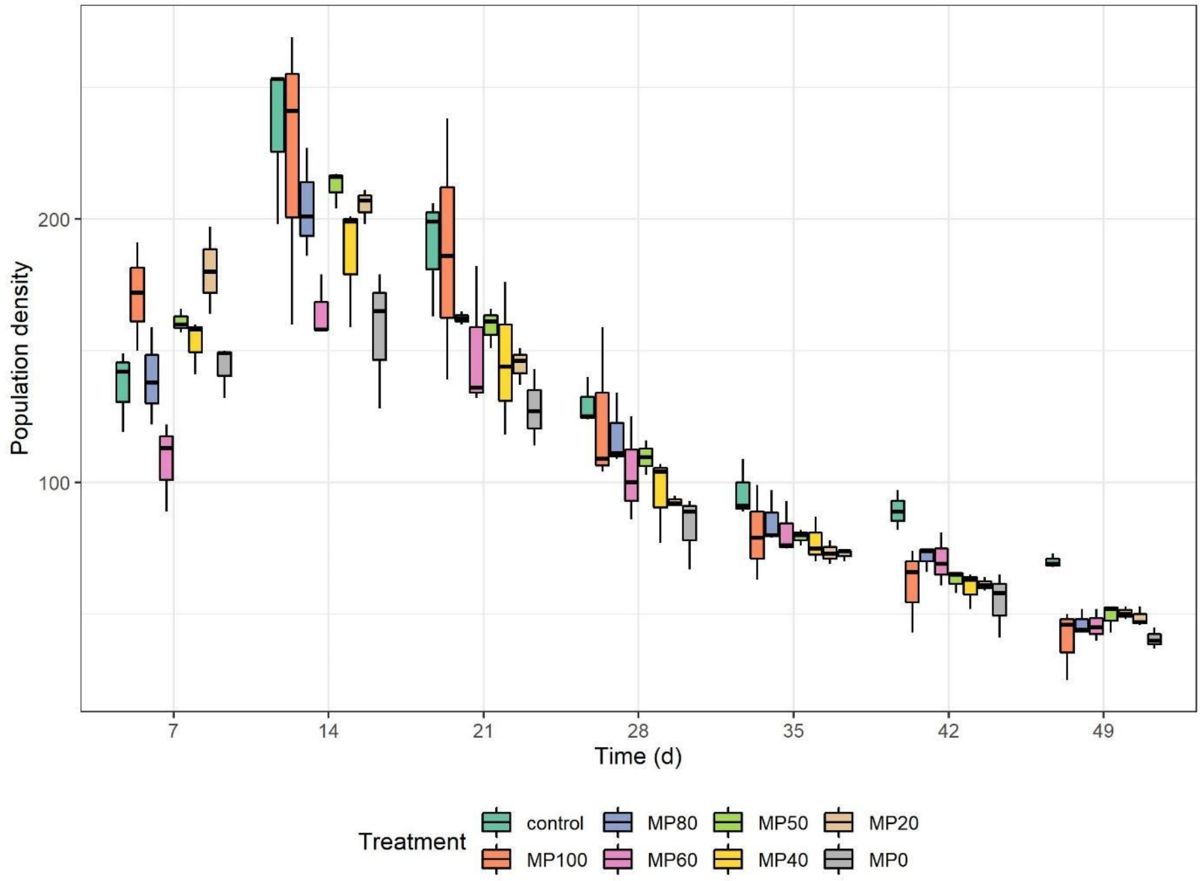

**Figure S4: Box-whisker plot of population density of *Daphnia magna* exposed to polystyrene microplastics (MP100), diatomite (MP0), or their mixtures over 50 d (n = 3).**

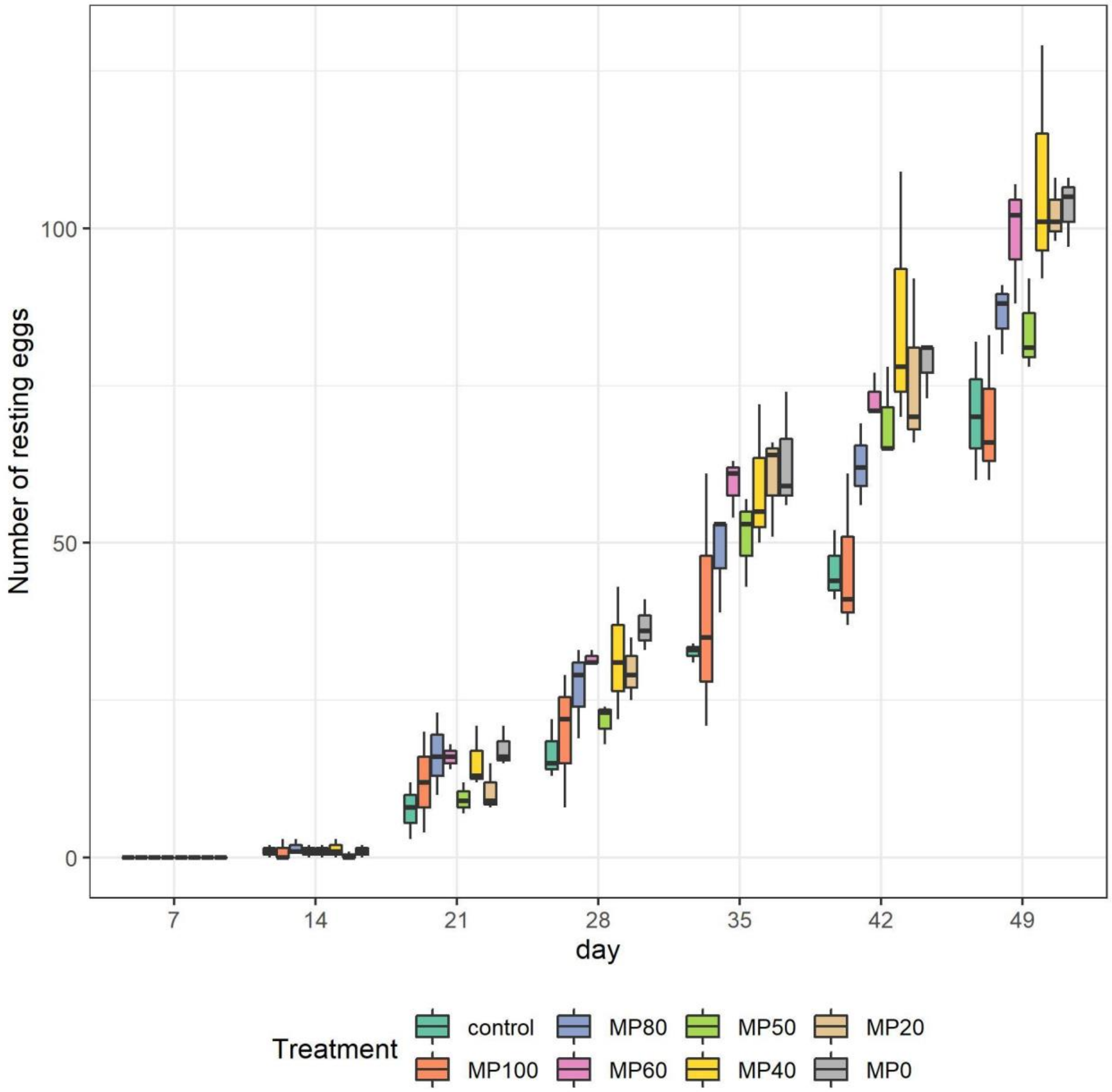

**Figure S5: Number of resting eggs produced by *Daphnia magna* populations exposed to polystyrene microplastics (MP100), diatomite (MP0), or their mixtures over 50 d (n=3).**

**Table S1: Results of the statistical comparison of the population size of *D. magna* individuals in populations exposed to particles compared to control populations.** One-way ANOVA with Holm-Šídák's multiple comparison tests for each observation time (adjusted p values).

| **Control vs.** | **d7** | **d14** | **d21** | **d28** | **d37** | **d42** | **d50** |
|---|---|---|---|---|---|---|---|
| **MP100** | 0.079 | 0.6232 | 0.9227 | 0.6845 | 0.1814 | **0.0079** | **0.0029** |
| **MP80** | 0.8383 | 0.5974 | 0.3956 | 0.6484 | 0.1866 | **0.0426** | **0.0031** |
| **MP60** | 0.1524 | **0.0423** | 0.2467 | 0.2732 | 0.1814 | **0.0426** | **0.0029** |
| **MP50** | 0.232 | 0.5974 | 0.3956 | 0.5013 | 0.1814 | **0.0083** | **0.0049** |
| **MP40** | 0.4994 | 0.2199 | 0.2243 | 0.1253 | 0.1408 | **0.0071** | **0.0049** |
| **MP20** | **0.0207** | 0.5974 | 0.2243 | 0.0938 | 0.0629 | **0.0079** | **0.0049** |
| **MP0** | 0.8383 | **0.0244** | 0.0535 | **0.0267** | 0.0617 | **0.0019** | **0.0006** |

**Table S2: Results of the statistical comparison of the number of *D. magna* neonates, juveniles and adults in populations exposed to particles compared to control populations.** One-way ANOVA with Holm-Šídák's multiple comparison tests for each size class and observation time (adjusted p values).

| **Control vs.** | **d7** | **d14** | **d21** | **d28** | **d37** | **d42** | **d50** |
|---|---|---|---|---|---|---|---|
| **Neonates** | | | | | | | |
| MP100 | 0.5321 | 0.4257 | 0.5161 | 0.1588 | 0.9623 | 0.2818 | 0.3224 |
| MP80 | 0.9864 | 0.3768 | 0.6793 | **0.0468** | 0.9623 | 0.2202 | 0.2403 |
| MP60 | 0.439 | **0.034** | 0.2713 | **0.0406** | 0.672 | 0.2202 | 0.1696 |
| MP50 | 0.9864 | 0.4257 | 0.5161 | 0.1588 | 0.7856 | 0.2202 | 0.4057 |
| MP40 | 0.6575 | 0.0681 | 0.2713 | **0.0406** | 0.9623 | 0.2202 | 0.4733 |
| MP20 | 0.8795 | 0.376 | 0.2713 | **0.0406** | 0.8379 | 0.1902 | 0.8349 |
| MP0 | 0.9864 | **0.0303** | 0.2686 | **0.0254** | 0.9623 | 0.2202 | 0.8349 |
| **Juveniles** | | | | | | | |
| MP100 | 0.9878 | 0.4237 | 0.983 | 0.9713 | 0.2534 | 0.1122 | **0.0004** |
| MP80 | 0.9878 | 0.9353 | 0.9496 | 0.9713 | 0.3383 | 0.3851 | **0.0028** |
| MP60 | 0.9878 | 0.974 | 0.983 | 0.9713 | 0.3383 | 0.3851 | **0.0005** |
| MP50 | 0.3845 | 0.9336 | 0.983 | 0.5914 | 0.3383 | 0.1122 | **0.0028** |
| MP40 | 0.9719 | 0.2876 | 0.983 | 0.9383 | 0.2534 | 0.0905 | **0.0017** |
| MP20 | 0.1298 | 0.6816 | 0.983 | 0.9174 | **0.0239** | 0.06 | **0.0005** |
| MP0 | 0.9878 | 0.974 | 0.983 | 0.5914 | **0.0079** | **0.0303** | **0.0008** |
| **Adults** | | | | | | | |
| MP100 | 0.3302 | 0.8893 | 0.8835 | 0.9936 | 0.9279 | 0.8901 | 0.9163 |
| MP80 | 0.5995 | 0.9465 | 0.5971 | 0.9936 | 0.9279 | 0.996 | >0.9999 |
| MP60 | 0.0807 | 0.3066 | 0.8214 | 0.9936 | 0.9432 | 0.8901 | 0.3774 |
| MP50 | 0.8224 | 0.9451 | 0.8214 | 0.2154 | 0.9432 | 0.996 | 0.9163 |
| MP40 | 0.8224 | 0.9465 | 0.7214 | 0.9936 | 0.9279 | 0.996 | 0.5598 |
| MP20 | 0.2052 | 0.9465 | 0.9886 | 0.9936 | 0.8417 | 0.9382 | 0.4414 |
| MP0 | 0.7363 | 0.9465 | >0.9999 | 0.8405 | 0.7596 | 0.996 | 0.9163 |

**Table S3: Mean population density as number of individuals per treatment group over time.**

| Treatment group | Day | Mean population density ± standard deviation |
|---|---|---|
| **Control** | 7 | 137 ± 16 |
| | 14 | 235 ± 32 |
| | 21 | 190 ± 23 |
| | 28 | 130 ± 9 |
| | 37 | 96 ± 11 |
| | 42 | 89 ± 8 |
| | 50 | 70 ± 3 |
| **MP100** | 7 | 171 ± 21 |
| | 14 | 223 ± 57 |
| | 21 | 188 ± 50 |
| | 28 | 124 ± 30 |
| | 37 | 80 ± 18 |
| | 42 | 61 ± 16 |
| | 50 | 45 ± 16 |
| **MP80** | 7 | 140 ± 19 |
| | 14 | 205 ± 21 |
| | 21 | 162 ± 3 |
| | 28 | 118 ± 14 |
| | 37 | 85 ± 10 |
| | 42 | 72 ± 5 |
| | 50 | 47 ± 5 |
| **MP60** | 7 | 108 ± 17 |
| | 14 | 165 ± 12 |
| | 21 | 150 ± 28 |
| | 28 | 104 ± 20 |
| | 37 | 81 ± 10 |
| | 42 | 70 ± 10 |
| | 50 | 46 ± 6 |
| **MP50** | 7 | 162 ± 5 |
| | 14 | 212 ± 7 |
| | 21 | 159 ± 8 |
| | 28 | 110 ± 9 |
| | 37 | 79 ± 3 |
| | 42 | 63 ± 4 |
| | 50 | 49 ± 6 |
| **MP40** | 7 | 153 ± 11 |
| | 14 | 186 ± 24 |
| | 21 | 146 ± 29 |
| | 28 | 96 ± 17 |
| | 37 | 77 ± 9 |
| | 42 | 60 ± 7 |
| | 50 | 50 ± 3 |
| **MP20** | 7 | 180 ± 17 |

| | | |
|---|---|---|
| | 14 | 205 ± 7 |
| | 21 | 145 ± 7 |
| | 28 | 93 ± 2 |
| | 37 | 73 ± 5 |
| | 42 | 61 ± 3 |
| | 50 | 49 ± 4 |
| **MP0** | 7 | 144 ± 10 |
| | 14 | 157 ± 26 |
| | 21 | 128 ± 15 |
| | 28 | 83 ± 14 |
| | 37 | 73 ± 2 |
| | 42 | 55 ± 12 |
| | 50 | 41 ± 4 |

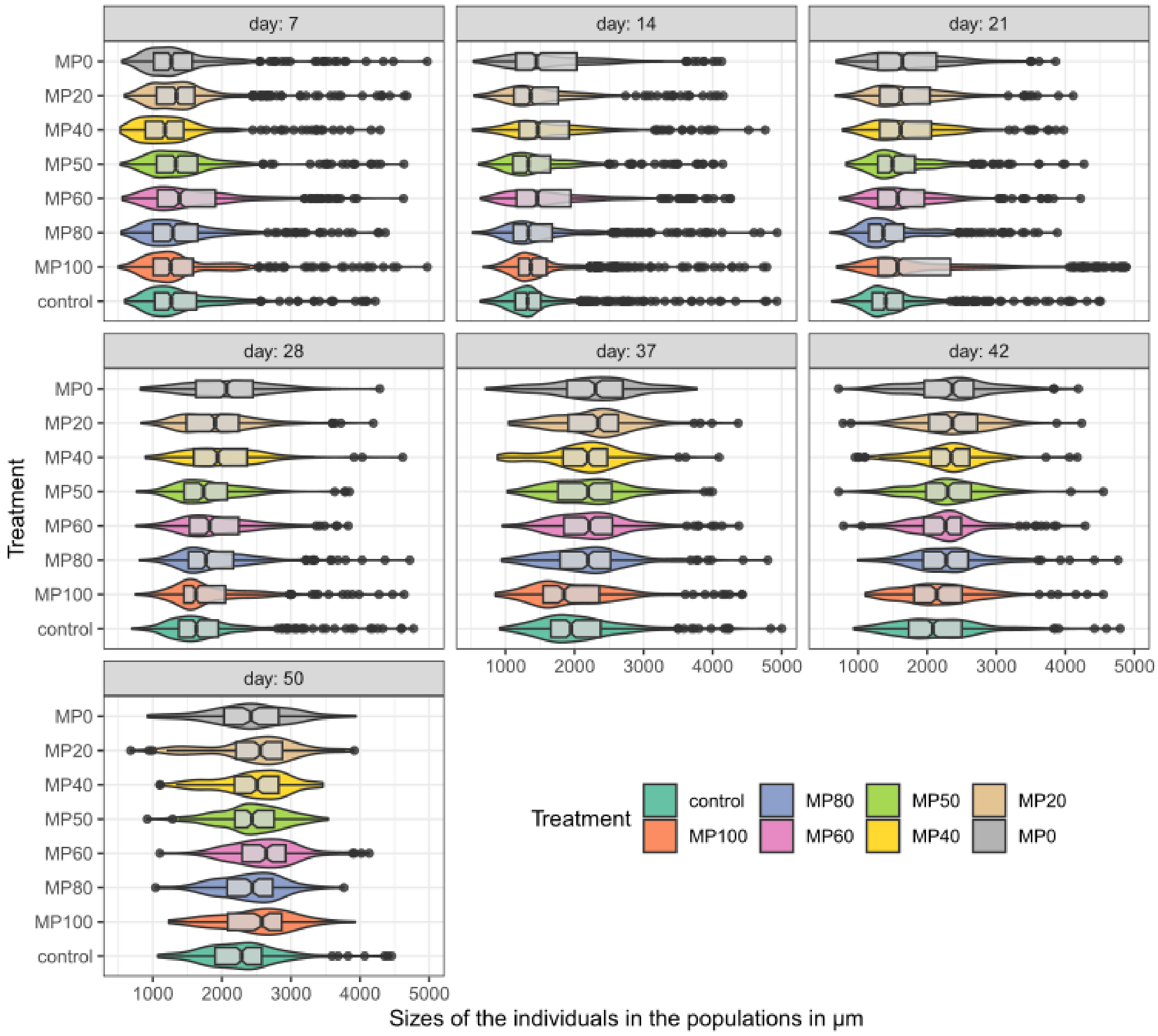

**Figure S6: Distributions of the body sizes of the individuals making up each treatment group per day.**

**Table S4: Abundance of age groups as number of neonates, juveniles, and adults per treatment over time.**

| Day | Treatment | Group | Mean abundance ± standard deviation |
|---|---|---|---|
| 7 | Control | Adults | 5.3 ± 2.1 |
| | | Juveniles | 46.3 ± 4.7 |
| | | Neonates | 85 ± 16.5 |
| | MP100 | Adults | 8.7 ± 2.3 |
| | | Juveniles | 50.7 ± 14 |
| | | Neonates | 111.7 ± 16.8 |
| | MP80 | Adults | 7.7 ± 1.5 |
| | | Juveniles | 48 ± 7.8 |
| | | Neonates | 84 ± 18.7 |
| | MP60 | Adults | 10.3 ± 3.2 |
| | | Juveniles | 42.7 ± 2.5 |
| | | Neonates | 55 ± 18.3 |
| | MP50 | Adults | 6.3 ± 1.2 |
| | | Juveniles | 65 ± 7.8 |
| | | Neonates | 89.7 ± 11 |
| | MP40 | Adults | 6.3 ± 1.5 |
| | | Juveniles | 39.7 ± 19.7 |
| | | Neonates | 107 ± 20.7 |
| | MP20 | Adults | 9.3 ± 1.5 |
| | | Juveniles | 72 ± 19 |
| | | Neonates | 99 ± 34 |
| | MP0 | Adults | 7 ± 3 |
| | | Juveniles | 46.7 ± 9.1 |
| | | Neonates | 90 ± 11.8 |
| 14 | Control | Adults | 9.7 ± 2.1 |
| | | Juveniles | 75.3 ± 6.4 |
| | | Neonates | 149.7 ± 37.4 |
| | MP100 | Adults | 12 ± 2 |
| | | Juveniles | 90 ± 12.8 |
| | | Neonates | 121.3 ± 45.8 |
| | MP80 | Adults | 11 ± 1.7 |
| | | Juveniles | 79.7 ± 7.6 |
| | | Neonates | 114 ± 13.5 |
| | MP60 | Adults | 15 ± 5.6 |
| | | Juveniles | 74.7 ± 7.8 |
| | | Neonates | 75.3 ± 16.3 |
| | MP50 | Adults | 7.7 ± 3.8 |
| | | Juveniles | 81 ± 6.1 |
| | | Neonates | 123.7 ± 7.6 |
| | MP40 | Adults | 8.3 ± 1.5 |
| | | Juveniles | 92.7 ± 12.1 |
| | | Neonates | 85.3 ± 32.1 |
| | MP20 | Adults | 9 ± 1.7 |
| | | Juveniles | 86 ± 11.8 |

| | | | |
|---|---|---|---|
| | | Neonates | 110.3 ± 16.6 |
| | MP0 | Adults | 11.3 ± 3.5 |
| | | Juveniles | 73.7 ± 11.7 |
| | | Neonates | 72.3 ± 34.9 |
| 21 | Control | Adults | 10.3 ± 3.1 |
| | | Juveniles | 83.3 ± 15 |
| | | Neonates | 95.7 ± 36.1 |
| | MP100 | Adults | 37 ± 49.4 |
| | | Juveniles | 85 ± 38.2 |
| | | Neonates | 98.5 ± 13.4 |
| | MP80 | Adults | 6.3 ± 0.6 |
| | | Juveniles | 70 ± 13.5 |
| | | Neonates | 86 ± 16 |
| | MP60 | Adults | 7.7 ± 1.2 |
| | | Juveniles | 91.7 ± 12.2 |
| | | Neonates | 50.7 ± 16.8 |
| | MP50 | Adults | 7.7 ± 3.8 |
| | | Juveniles | 86.3 ± 4 |
| | | Neonates | 65.3 ± 4.2 |
| | MP40 | Adults | 7 ± 3.6 |
| | | Juveniles | 92.3 ± 11.9 |
| | | Neonates | 46.7 ± 30.4 |
| | MP20 | Adults | 10 ± 5.3 |
| | | Juveniles | 88.7 ± 9.3 |
| | | Neonates | 46 ± 18.5 |
| | MP0 | Adults | 10.3 ± 0.6 |
| | | Juveniles | 73.7 ± 4 |
| | | Neonates | 44 ± 16.7 |
| 28 | Control | Adults | 12 ± 3.5 |
| | | Juveniles | 83.7 ± 2.5 |
| | | Neonates | 34 ± 9.2 |
| | MP100 | Adults | 11.3 ± 2.1 |
| | | Juveniles | 89.3 ± 24 |
| | | Neonates | 23.3 ± 8.5 |
| | MP80 | Adults | 11 ± 6 |
| | | Juveniles | 90.7 ± 13.4 |
| | | Neonates | 16.3 ± 3.1 |
| | MP60 | Adults | 11 ± 4.4 |
| | | Juveniles | 78 ± 21.3 |
| | | Neonates | 14.7 ± 4.5 |
| | MP50 | Adults | 6 ± 1.4 |
| | | Juveniles | 83 ± 2.8 |
| | | Neonates | 20.5 ± 13.4 |
| | MP40 | Adults | 10.7 ± 4.6 |
| | | Juveniles | 71.7 ± 4.7 |
| | | Neonates | 13.7 ± 12.5 |
| | MP20 | Adults | 10.3 ± 5 |
| | | Juveniles | 68.3 ± 6.7 |
| | | Neonates | 14 ± 3 |

| | | | |
|---|---|---|---|
| | MP0 | Adults | 16.3 ± 3.2 |
| | | Juveniles | 55 ± 8.5 |
| | | Neonates | 11.7 ± 5.8 |
| 37 | Control | Adults | 16 ± 1 |
| | | Juveniles | 69.7 ± 6.7 |
| | | Neonates | 10.7 ± 9 |
| | MP100 | Adults | 12 ± 2.6 |
| | | Juveniles | 56.3 ± 15.7 |
| | | Neonates | 12 ± 7.2 |
| | MP80 | Adults | 18.3 ± 8.1 |
| | | Juveniles | 59.3 ± 8.1 |
| | | Neonates | 7.7 ± 4 |
| | MP60 | Adults | 17.3 ± 2.5 |
| | | Juveniles | 61 ± 9.6 |
| | | Neonates | 3 ± 1.7 |
| | MP50 | Adults | 16 ± 2.6 |
| | | Juveniles | 59 ± 1.7 |
| | | Neonates | 4.3 ± 4.9 |
| | MP40 | Adults | 12 ± 6.6 |
| | | Juveniles | 56 ± 4.6 |
| | | Neonates | 9.3 ± 9.7 |
| | MP20 | Adults | 20.7 ± 6 |
| | | Juveniles | 47.3 ± 4 |
| | | Neonates | 5.3 ± 4 |
| | MP0 | Adults | 21.7 ± 7.4 |
| | | Juveniles | 43.3 ± 6.5 |
| | | Neonates | 7.7 ± 4 |
| 42 | Control | Adults | 17.3 ± 6.7 |
| | | Juveniles | 65 ± 10.8 |
| | | Neonates | 7 ± 3 |
| | MP100 | Adults | 12.7 ± 6.4 |
| | | Juveniles | 43.7 ± 19.6 |
| | | Neonates | 4.7 ± 3.2 |
| | MP80 | Adults | 16 ± 2.6 |
| | | Juveniles | 53.3 ± 6 |
| | | Neonates | 2.3 ± 1.5 |
| | MP60 | Adults | 13 ± 5.6 |
| | | Juveniles | 54 ± 15.7 |
| | | Neonates | 3.3 ± 3.2 |
| | MP50 | Adults | 17 ± 3.6 |
| | | Juveniles | 43.3 ± 1.5 |
| | | Neonates | 2.7 ± 0.6 |
| | MP40 | Adults | 16.3 ± 4.7 |
| | | Juveniles | 41.3 ± 8.6 |
| | | Neonates | 3.5 ± 2.1 |
| | MP20 | Adults | 20.7 ± 4.9 |
| | | Juveniles | 38.7 ± 4.6 |
| | | Neonates | 3 ± 2.8 |
| | MP0 | Adults | 16.7 ± 3.8 |

| | | | |
|---|---|---|---|
| | | Juveniles | 35 ± 10.1 |
| | | Neonates | 4.5 ± 0.7 |
| 50 | Control | Adults | 17 ± 1.7 |
| | | Juveniles | 49.7 ± 4.5 |
| | | Neonates | 3.3 ± 1.5 |
| | MP100 | Adults | 19.3 ± 2.3 |
| | | Juveniles | 20 ± 12.1 |
| | | Neonates | 1.5 ± 0.7 |
| | MP80 | Adults | 17 ± 3.6 |
| | | Juveniles | 28.7 ± 7 |
| | | Neonates | 2 ± NA |
| | MP60 | Adults | 24 ± 6.1 |
| | | Juveniles | 21.3 ± 1.5 |
| | | Neonates | 1 ± NA |
| | MP50 | Adults | 19 ± 4 |
| | | Juveniles | 29 ± 8.7 |
| | | Neonates | 2 ± 1.4 |
| | MP40 | Adults | 22.3 ± 3.1 |
| | | Juveniles | 26.3 ± 5.8 |
| | | Neonates | 1.7 ± 0.6 |
| | MP20 | Adults | 23.3 ± 6.4 |
| | | Juveniles | 21.3 ± 4.9 |
| | | Neonates | 4 ± 2 |
| | MP0 | Adults | 14.3 ± 5 |
| | | Juveniles | 23.7 ± 2.1 |
| | | Neonates | 4 ± 1.4 |

**Table S5: Number of resting eggs per replicate after 50 d of exposure.**

| Treatment group | Replicate | Resting eggs |
|---|---|---|
| Control | 1 | 82 |
| | 2 | 70 |
| | 3 | 60 |
| MP100 | 1 | 83 |
| | 2 | 66 |
| | 3 | 60 |
| MP80 | 1 | 88 |
| | 2 | 91 |
| | 3 | 80 |
| MP60 | 1 | 102 |
| | 2 | 107 |
| | 3 | 88 |
| MP50 | 1 | 78 |
| | 2 | 81 |
| | 3 | 92 |
| MP40 | 1 | 129 |
| | 2 | 101 |
| | 3 | 92 |
| MP20 | 1 | 108 |
| | 2 | 101 |
| | 3 | 98 |
| MP0 | 1 | 105 |
| | 2 | 97 |
| | 3 | 108 |

**Table S6: Results of the statistical comparison of the body length of *D. magna* individuals in populations exposed to particles.** Kruskal-Wallis tests with Dunn's multiple comparison tests for each observation time. Δ rank indicates the difference in mean rank (negative values imply larger individuals) and p refers to the adjusted p values.

| | **d 7** | | **d 14** | | **d 21** | | **d 28** | | **d 37** | | **d 42** | | **d 50** | |
|---|---|---|---|---|---|---|---|---|---|---|---|---|---|---|
| **Control vs.** | **Δ rank** | **p** | **Δ rank** | **p** | **Δ rank** | **p** | **Δ rank** | **p** | **Δ rank** | **p** | **Δ rank** | **p** | **Δ rank** | **p** |
| MP100 | 46.5 | >0.9999 | -271.5 | **0.0018** | -26 | >0.9999 | -67.9 | >0.9999 | 65.9 | >0.9999 | -33.9 | >0.9999 | -156.5 | **0.0004** |
| MP80 | 2 | >0.9999 | -150 | 0.3391 | 68.6 | >0.9999 | -210.5 | **0.0004** | -145.8 | **0.0168** | -139.7 | **0.0064** | -90.4 | 0.1029 |
| MP60 | -223.2 | **0.0259** | -452 | **<0.0001** | -363.9 | **<0.0001** | -279.6 | **<0.0001** | -193 | **0.0005** | -101.9 | 0.1129 | -221.6 | **<0.0001** |
| MP50 | -86.6 | >0.9999 | -87.5 | >0.9999 | -243.9 | **0.0014** | -133.7 | 0.1807 | -142.4 | **0.0255** | -148 | **0.005** | -112.1 | **0.0144** |
| MP40 | 322 | **<0.0001** | -469 | **<0.0001** | -487.8 | **<0.0001** | -383.6 | **<0.0001** | -115.9 | 0.1315 | -175.6 | **0.0005** | -136.6 | **0.0011** |
| MP20 | -91.5 | >0.9999 | -196.1 | 0.0686 | -475.6 | **<0.0001** | -298.7 | **<0.0001** | -260.8 | **<0.0001** | -210.7 | **<0.0001** | -157.5 | **0.0001** |
| MP0 | 21 | >0.9999 | -442 | **<0.0001** | -446.1 | **<0.0001** | -470.2 | **<0.0001** | -255.5 | **<0.0001** | -183.3 | **0.0004** | -94.19 | 0.1027 |
